# Improved full-length killer cell immunoglobulin-like receptor transcript discovery in Mauritian cynomolgus macaques

**DOI:** 10.1101/093559

**Authors:** Trent M. Prall, Michael E. Graham, Julie A. Karl, Roger W. Wiseman, Adam J. Ericsen, Muthuswamy Raveendran, R. Alan Harris, Donna M. Muzny, Richard A. Gibbs, Jeffrey Rogers, David H. O’Connor

## Abstract

Killer cell Immunoglobulin-like Receptors (KIRs) modulate disease progression of pathogens including HIV, malaria, and hepatitis C. Cynomolgus and rhesus macaques are widely used as nonhuman primate models to study human pathogens and so considerable effort has been put into characterizing their KIR genetics. However, previous studies have relied on cDNA cloning and Sanger sequencing that lacks the throughput of current sequencing platforms. In this study, we present a high throughput, full-length allele discovery method utilizing PacBio circular consensus sequencing (CCS). We also describe a new approach to Macaque Exome Sequencing (MES) and the development of the Rhexome1.0, an adapted target capture reagent that includes macaque-specific capture probesets. By using sequence reads generated by whole genome sequencing (WGS) and MES to inform primer design, we were able to increase the sensitivity of KIR allele discovery. We demonstrate this increased sensitivity by defining nine novel alleles within a cohort of Mauritian cynomolgus macaques (MCM), a geographically isolated population with restricted KIR genetics that was thought to be completely characterized. Finally, we describe an approach to genotyping KIRs directly from sequence reads generated using WGS/MES reads. The findings presented here expand our understanding of KIR genetics in MCM by associating new genes with all eight KIR haplotypes and demonstrating the existence of at least one *KIR3DS* gene associated with every haplotype.

## INTRODUCTION

Killer cell Immunoglobulin-like receptors (KIRs) are a complex family of receptors expressed on the surface of natural killer (NK) cells and subsets of T lymphocytes (Lodoen and Lanier 2006). KIRs modulate immune responses through interactions with major histocompatibility complex (MHC) class I molecules expressed on target cell surfaces (Lanier 2005; Sambrook et al. 2006). KIRs can possess either two Ig-like domains (KIR2D) or three Ig-like domains (KIR3D), a transmembrane domain and either long (L) or short (S) cytoplasmic tails capable of eliciting inhibitory or activating signaling cascades, respectively (Gardiner 2008). KIRs with one Ig-like domain and a truncated cytoplasmic tail (KIR1D) are present in some nonhuman primates, but are absent in humans (Bimber and Evans 2015; Hershberger et al. 2005).

Human KIR genes are located within the leukocyte receptor complex (LRC) on chromosome 19q14.3 along with other immunoglobulin superfamily receptor genes (Trowsdale et al. 2001; Wende et al. 2000). This genomic region contains anywhere from seven to twelve KIR genes. Fifteen genes (and two pseudogenes) have been characterized in humans to date (Carrington and Norman 2003; Uhrberg et al. 1997). Each chromosome contains a group of KIR genes that are inherited together; collectively, these comprise a KIR haplotype. The combined genes of both haplotypes define an individual’s KIR genotype. Human KIR haplotypes are defined by the presence or absence of genes other than *KIR3DL3, KIR3DP1, KIR2DL4,* and *KIR3DL2,* which are found on all haplotypes and are considered framework genes (Wilson et al. 2000). Haplotypes can be broadly categorized into group A haplotypes and more complex group B haplotypes. Group A haplotypes always contain seven loci which include the framework genes and a single activating receptor, *KIR2DS4.* Group B haplotypes contain variable numbers of genes that include multiple activating receptors other that *KIR2DS4.* Each KIR gene exhibits a high degree of nucleotide sequence polymorphism (Middleton and Gonzelez 2010). As of February 2015 (release 2.6.1), 753 unique, full-length human KIR alleles have been characterized and catalogued for humans in the Immuno Polymorphism Database (Robinson et al. 2005).

Each KIR binds a specific set of ligands and so the genes encoded within an individual’s genome determine, in part, the specificity of potential immune responses (Carrillo-Bustamante et al. 2016; Long and Rajagopalan 2000). NK cells only express a subset of an individual’s KIR genes. However, all functional KIR genes within an individual’s genome are expressed by at least one subpopulation of NK cells (Gardiner 2008; Valiante et al. 1997). Sequence polymorphisms of KIR genes are known to play an important role in levels of cell surface expression as well as binding affinities and NK cell effector function (Biassoni et al. 1995; Carr et al. 2005; Dunphy et al. 2015; Hilton et al. 2015). For instance, Yawata et al. (2006) showed that *KIR3DL1*001*,*KIR3DL1 *020*, and *KIR3DL1*01502* alleles exhibited higher antibody binding and were expressed by larger proportions of NK cells than *KIR3DL1 *005* and *KIR3DL1 *007* alleles. Furthermore, it was shown that the inhibitory capacity of NK cells was greater for high-expressing *KIR3DL1* alleles save for KIR3DL1*005. Sequence polymorphisms between HLA genes can also influence the relative abundance of NK cell populations resulting in unequal representation of KIRs within the repertoire (Shilling et al. 2002; Yawata et al 2006). Taken together, compound KIR/HLA genotypes are crucial in the determination of an individual’s receptor repertoire and signaling capabilities.

Compound KIR/HLA genotypes have been associated with protection from a wide spectrum of pathogens including HIV, hepatitis C, malaria, and papilloma virus (Alter et al. 2007; Bonagura et al. 2010; Hansen et al. 2007; Khakoo et al. 2004; Knapp et al. 2010; Zipperlen et al. 2015). The role of *KIR3DS1* and *HLA-Bw4-80* Ile has been evaluated in the context of HIV infection. Multiple groups have observed an association between *KIR3DS1+ and HLA-Bw4-80Ile+* genotypes and slower progression to AIDS (Jiang et al. 2013; Martin et al. 2002). Functionally, *KIR3DS1+* NK control HIV replication within autologous CD4+ T cells *in vitro* (Alter et al. 2007; Zipperlen et al. 2015). KIR genotypes also affect transplantation outcomes. Donor/recipient KIR ligand mismatches reduce graft survival in hepatic and renal transplantation (Kuśnierczyk 2013; La Manna et al. 2013; Legaz et al. 2013; Van Bergen et al. 2011). For instance, Legaz et al. and Cirocco et al. independently found that acute rejection was significantly decreased in transplants involving *KIR2DL2+* recipients matched with *HLA-C1+* donors compared to *HLA-C1-* donors (Cirocco et al. 2007; Kunert et al. 2007). Although our understanding of KIR signaling has significantly advanced in recent years, the mechanisms governing these differential immune responses remains elusive due to the complexity of KIR/HLA genetics.

Cynomolgus and rhesus macaques *(Macaca fascicularis, Macaca mulatta)* are widely used model organisms in the study of human infectious diseases and transplantation due to their genetic homology and ability to mount similar immune responses (Anderson and Kirk 2013; Antony and MacDonald 2015; Messaoudi et al. 2011; Schmitz and Korioth-Schmitz 2013). Significant effort has been put into characterizing the genetic diversity of KIRs and MHC in macaques, particularly in rhesus macaques of Indian origin (Blokhuis et al. 2010; Blokhuis et al. 2011; Hershberger et al. 2001; Kruse et al. 2010; Moreland et al. 2011; Sambrook et al. 2005; Wiseman et al. 2009). Twenty-two KIR genes (referred to as *Mamu-KIR)* are thought to exist in Indian rhesus macaques and a total of 149 full-length sequence variants of these genes are currently deposited in GenBank (Bimber and Evans 2015). The average number of expressed genes per animal is twelve (Blokhuis et al. 2011). Kruse et al. (2010) found five to eleven KIR genes/haplotype in their Indian rhesus families. In this study, they concluded, “rhesus macaque and human KIR haplotypes show a comparable level of diversity and complexity.” A study by Blokhuis et al. (2011) found 272 unique genotypes within a relatively small population of 378 rhesus macaques. For comparison, only 573 KIR genotypes have been characterized in humans despite screening 18,783 individuals from 155 different populations to date (González-Galarza et al. 2015). It is therefore likely that the diversity of KIR genotypes in macaques is at least comparable to, and may even exceed, that of humans. Because studying KIRs requires genetic control over both KIR and MHC genotypes, the extreme diversity of *Mamu-KIR* genotypes makes establishing a cohort of KIR/MHC identical rhesus macaques virtually impossible.

Mauritian cynomolgus macaques (MCM) provide a possible alternative due to their restricted genetic diversity. MCM are descended from a small founder population introduced on the island of Mauritius approximately 500 years ago (Sussman et al. 1986). This founder effect can be seen through MHC genetics in which seven haplotypes account for virtually all variation within the population (Wiseman et al. 2013). In a previous study we characterized the genetic diversity of KIRs in MCM (referred to as *Mafa-KIR)* using microsatellite analysis within a cohort of 274 MCM. Similar to MHC, eight KIR haplotypes were sufficient in describing essentially all genetic variation within the MCM population (Bimber et al. 2008). Cloning and Sanger-based sequencing identified a total of forty alleles and eleven splice variants that segregated into one *KIR1D*, one *KIR3DS*, two *KIR2DL*, and four presumptive *KIR3DL* lineages.

Previous efforts to characterize KIRs in macaques have utilized cloning and Sanger sequencing, allele lineage-specific PCR, microsatellite analysis, and Roche/454 pyrosequencing (Bimber et al 2008; Blokhuis et al. 2010; Blokhuis et al. 2011; Kruse et al. 2010; Moreland et al. 2011; Sambrook et al. 2005). Although informative, these methods are time and resource intensive or cannot resolve allele-level genotypes unequivocally due to short products. Pacific Biosciences (PacBio) circular consensus sequencing (CCS) with single-molecule real-time (SMRT) technology produces long reads with high redundancy (Eid et al. 2009, Travers et al. 2010; Westbrook et al. 2015). Long reads eliminate the uncertainty of short read assembly required by previous massively parallel sequencing approaches to resolve full-length KIR transcripts. Here we present a novel full-length KIR allele discovery method utilizing the PacBio RSII platform. We increased the sensitivity of our assay by utilizing whole genome and macaque exome sequencing (WGS/MES) data to design primers in conserved regions of the genome. We then constructed a *Mafa-KIR* allele library based on PacBio amplicon sequences and demonstrated *Mafa-KIR* genotypes can be derived directly from genomic sequence reads. This study provides a new understanding of *Mafa-KIR* genetics and highlights the power of applying long-read deep sequencing techniques to KIRs and other complex immune loci.

## MATERIALS AND METHODS

### Animal Selection

Blood samples from 30 MCM were selected for KIR analysis (Supplementary Table 1). KIR genotypes for 10 animals were determined through microsatellite analysis in a previous study representing at least one chromosome with a non-recombinant KIR region for each of the eight KIR haplotypes in the MCM population (Bimber et al. 2008).

### cDNA amplicons and PacBio RS II Sequencing

RNA was isolated from peripheral blood mononuclear cells (PBMCs) or whole blood using the Promega Maxwell 16 MDx Instrument and Maxwell 16 LEV simplyRNA Cells or Maxwell 16 LEV simplyRNA Blood kits (Promega, Madison, WI, USA) according to manufacturer’s protocols. First strand complementary DNA (cDNA) was synthesized using SuperScript III First-Strand Synthesis System (Invitrogen, Carlsbad, CA, USA).

To amplify complete KIR open reading frames (ORFs), cDNA-PCR was performed to create amplicons ranging from 500-1600 bp using Phusion high-fidelity polymerase (New England Biolabs, Ipswich, MA, USA). The following primer sequences were used to generate templates for sequencing: KIR1D/3D_F (5’-CTKTCTGMACCGGCAGCACC-3’), KIR1D/3D_R (5’-GGGGTCAAGTGAAGTGGAGA-3’), KIR2DL4_F (5’-GAGTCACTGCATCCTGGCA-3’), KIR2DL4_R (5’-CGCAGACGTTGGTAAGCAAG-3’). Independent reactions were performed to generate amplicons for *KIR1D/3D* and *KIR2DL4* products and the resulting products for each animal were pooled prior to PacBio library construction. A unique 16-bp barcode was appended to the 5’ end of each primer from a set of 384 barcodes designed for the PacBio system. Each sample could then be identified by its unique combination of barcodes. For all reactions, thermal cycling conditions were: denaturation at 98°C for 30 s; 30 cycles of 98°C for 5 s; 64°C *(KIR3DL/KIR3DS/KIR1D)* or 56°C *(KIR2DL4)* for 10 s, 72°C for 20 s; and a final extension of 72°C for 5 min. Successful amplification was confirmed using the FlashGel System (Lonza Group Ltd., Basel, Switzerland).

Amplified DNA was purified twice using Ampure XP SPRI beads (Agencourt Bioscience Corporation, Beverly, MA, USA) at a 0.6:1 bead to DNA volume ratio. All samples were quantified using the Qubit dsDNA BR assay kit and Qubit 2.0 Fluorometer (Thermo Fisher Scientific, Waltham, MA, USA). Amplicon size distributions were measured using the Agilent 2100 Bioanalyzer DNA12000 kit (Life Technologies, Madison, WI, USA). SMRTbell libraries were created using the PacBio Amplicon Template Preparation protocol for CCS (http://www.pacb.com). This protocol allows individual molecules to be sequenced multiple times in both orientations. Briefly, a pool of primary PCR products was end-repaired and hairpin adapters were incorporated using the PacBio DNA Template Prep Kit 2.0. After removal of failed ligation products using exonuclease III and VII, SMRTbells were purified using a 0.6:1 volume ratio of AMPure PB beads to DNA. Concentrations and sizes were obtained again for the final library prep using the Qubit dsDNA BR assay and Agilent 2100 Bioanalyzer DNA12000 kits. The volume of sequencing primer and polymerase was determined using a PacBio calculator by entering the concentrations of SMRTbells and ~1200 bp average insert size. Sequencing primer was annealed to the single-stranded loop region of the SMRTbell template and primer-annealed SMRTbells were bound to DNA polymerase P6. Polymerase-bound SMRTbells were MagBead loaded over zero-mode waveguides and immobilized. Sequencing was performed by the Great Lakes Genomics core at the University of Wisconsin-Milwaukee using a PacBio RS II instrument with C4 sequencing chemistry. PacBio CCS generates long read lengths (>10–15 kb) from a single molecule allowing unambiguous phasing of alleles.

### Pacific Biosciences Data Analysis

The Pacific Biosciences command line tool pbccs from the smrttools package version 3.0.1 (https://github.com/PacificBiosciences/pbccs) was used to produce CCS from raw data files. This tool produced amplicon sequences in the form of BAM files, which were not demultiplexed. These BAM files were then filtered on the Read Quality field or “rq” tag and sequences with an “rq” value of 0.99 or higher were saved. This data was used for the purpose of novel allele discovery. Pacific Biosciences provided tools pbbarcode and bamCCSBarcodeDemultiplex.py were used to demultiplex the read data by barcode. A barcode score threshold was set at 25 to accurately match barcode sequences within reads. Demultiplexed fastq files were used for genotyping individual samples.

A series of analysis steps with open-source bioinformatics utilities were performed to discover putative novel allele sequences within pre-processed Pacific Biosciences sequencing data. As a first step after read pre-processing, chimera filtering was performed using the USEARCH (Edgar 2010) software package and the uchime_ref algorithm with a reference KIR allele sequence database. Following chimera filtering, primers were trimmed from read ends using the bbduk software tool as part of the bbtools (https://sourceforge.net/projects/bbmap/files/) software package. Primers were trimmed from the left and right ends of sequencing reads sequentially, using bbduk’s parameters “ktrim=l” and “ktrim=r” to specify which end to trim. bbduk’s parameters “restrictleft=50” and “restrictright=50” were used to limit the region of a sequencing read considered for primer trimming and a kmer size of 16 was specified with “k=16”. Next, open reading frames with sizes between 400 and 1300 base pairs were identified within read sequences using the software tool getorf as part of the EMBOSS software package (Rice et al. 2000). Only sequence reads containing an intact ORF within the specified size range were included in further processing steps. Reads identical to previously described allele sequences were then removed by mapping them to a database of known KIR allele sequences using mapPacBio.sh as part of the bbtools software package, and requiring reads match a reference allele sequence perfectly end to end using the parameter “perfectmode=t”. Unmapped reads were then clustered using the cluster_fast algorithm within the USEARCH software package, requiring 100% identity between sequences in a cluster. Cluster representative sequences with 3 or more reads in support of the cluster were saved for further analysis.

Putative novel allele sequences having been filtered by the previously mentioned analysis steps were then imported into the software tool Geneious Pro R9 (http://www.geneious.com, Kearse et al., 2012 http://www.geneious.com, Kearse et al., 2012). Circular consensus sequences were mapped to the set of putative novel allele sequences within Geneious (requiring 100% identity in the overlap) using the bbmap plugin and parameters “semiperfectmode” and “PacBio mode”’. Putative novel sequences meeting the requirement of having greater than or equal to 3 reads whose mapping spanned the complete length of the sequence, and had at least 3 reads from a single specific sample were considered valid novel sequences and included in genotyping analyses.

To genotype the samples within the sequencing run, the set of valid novel sequences was merged with the set of previously described reference allele sequences. Barcode binned circular consensus sequences were then mapped to this merged set of allele sequences using Geneious Pro R9 (requiring 100% identity in the overlap). The mapping results were then exported from Geneious Pro R9 and imported into Microsoft Excel 2011 to generate a table of genotypes.

### Whole Genome Sequencing (WGS)

WGS datasets from 18 MCM were generated in a previous study (Supplementary Table 2) (Ericsen et al. 2014). Briefly, genomic DNA from was isolated from PBMCs and Illumina paired-end libraries were constructed following standard protocol (complete description available at http://www.hgsc.bcm.edu). Library templates were prepared with TruSeq PE Cluster Generation Kits (Illumina, San Diego, CA, USA) using Illumina’s cBot cluster generation system. Libraries were loaded onto three HiSeq flow cell lanes and paired-end sequencing reads were generated using the Illumina HiSeq 2000 platform. Sequencing-by-synthesis reactions were extended for 101 cycles from each end using TruSeq SBS Kits (Illumina, San Diego, CA, USA). On average, 118 Gb (approximately 44Gb per lane) of sequence was generated per sample.

### Rhexome1.0 target capture reagent

To improve overall coverage of the rhesus exome and successfully sequence the macaque MHC region, capture probes were designed targeting rhesus exons that were not covered sufficiently using the human based NimbleGen VCRome2.1 capture reagents. Sequencing reads from rhesus macaque DNA samples captured using VCRome2.1 were mapped to the rheMac2 genome assembly. Macaque exons that did not pass read coverage thresholds were identified (exons that showed less than 20x coverage or had excessive coverage were considered failures). In addition, all exons plus introns and UTR sequences for each class I and class II loci from the human MHC region (hg19) were included in this custom probeset to increase the number of MHC class I and II sequence reads recovered in each experiment.

The resulting set of novel exonic and MHC targets sequences were provided to NimbleGen scientists (NimbleGen, Madison, WI, USA), where they used proprietary software to design and manufacture novel sequence capture probes. This sequence capture probe set (macaque exonic regions: 1,544,913bp; human MHC region: 607,629bp, hg19) was used as a “spike-in” reagent added to the VCRome2.1 probe set in standard human exome capture procedures (Lupski et al. 2013). A series of titrations comparing the results for exome sequencing using the combined VCRome2.1 plus “spike-in” reagents against whole genome sequencing results from the same animals showed that a ratio of 1:2.5 for VCRome2.1-to-“spike-in” provided superior overall coverage results. We have designated the resulting VCRome2.1 plus “spike-in” reagents the Rhexome1.0.

### Whole Exome Sequencing (WES) and Macaque Exome Sequencing (MES)

Eight MCM exomes using standard VCRome2.1 capture reagents and 36 MCM exomes using Rhexome1.0 capture reagents were prepared using standard human exome capture procedures (Supplementary Table 2) (Lupski et al. 2013). After standard DNA quality control tests on samples, we produced Illumina sequencing libraries with incorporated barcodes, following standard procedures (Lupski et al. 2013). Groups of six macaque bar-coded samples were pooled and captured together (6Plex). Each resulting pool of six DNA samples (enriched for the macaque exome by the capture process) was sequenced in one lane of Illumina HiSeq2000. This procedure results in an estimated sequence read depth of greater than 20X for 91% of on-target reads. For distinction, whole exome sequencing (WES) describes sequencing experiments that utilize standard VCRome2.1 whereas macaque exome sequencing (MES) describes experiments that utilize Rhexome1.0.

### Genotyping from WGS and MES datasets

A reference allele sequence database was created consisting of discriminatory exon regions found within the complete set of full-length reference alleles having been discovered from PacBio sequencing. For *KIR1D*, *KIR2DL4*, *KIR3DS* and *KIR3DL* loci, exon sets were chosen based on having polymorphisms that enabled allelic resolution. For *KIR1D*, D1 exon sequence was used for each full-length allele; for *KIR2DL4*, D0 and D2 exon sequences were used for each full-length allele; and for *KIR3DS/KIR3DL*, D0, D1, and D2 exon sequences were used for each full-length allele. The genotyping rationale used was that mapping results that included specific exon sequences for a given KIR locus being completely covered 5’ to 3’ with reads that map with 100% identity represents the presence of the associated full length allele sequence in the sample being genotyped.

A custom software pipeline was written in Python and used to run sequential analysis steps for genotyping samples from WGS/MES datasets (https://bitbucket.org/dholab/mcm_kir/src). As a first step, read sorting and read pairing verification was done using the repair.sh software tool within the bbtools (https://sourceforge.net/projects/bbmap/files/) software package. Next, reads were mapped to the exon allele reference database that was created using bbmap as part of the bbtools (https://sourceforge.net/projects/bbmap/files/) software package. Reads were required to map with 100% identity in the overlap with the exon reference sequences. Mapping results were then translated to coverage statistics using coverage Bed within the bedtools (Quinlan and Hall 2010) software package. Mapping results were then filtered using custom Python code based on the coverage statistics to only include results where exon reference sequences had at least 1 read covering every base of the reference. Filtering was also done to require the correct number of discriminatory exons be covered for each KIR locus. Mapping results were then aggregated and read counts broken out by exon for each sample per KIR allele sequence. The results were reported as a genotyping table.

### Phylogenetic Analysis

Codon based nucleotide alignments were generated in Geneious Pro R9 (Kearse et al. 2012) using the ClustalW2 plugin (Larkin et al. 2007) with default settings. jModelTest2 (Darriba et al. 2012; Guindon and Gascuel 2003) was used to statistically select GTR+I+γ as the best fit nucleotide substitution model from sequence alignments. Maximum likelihood trees were generated using the RaXML plugin (GTR+I+λ, model). Trees were refined using bootstrapping (1,000 replicates).

## RESULTS

### Full-length KIR allele discovery

Previously, we designed a primer pair to amplify full-length KIR transcripts based on rhesus macaque cDNA transcripts deposited in NCBI Genbank as of 2008. We adapted this primer pair for PacBio sequencing in a pilot study with a cohort of twelve MCM. As expected *KIR1D* and *KIR3D* sequences were recovered that had been characterized previously but we were unable to detect any *KIR2DL4* sequences. To investigate whether this reflected variation within the binding sites of primers for *KIR2DL4* transcripts, we compared our original primer sequences against genomic sequences from 62 total WGS, WES, and MES MCM samples (Ericsen et al. 2014). In order to distinguish KIR2DL4-specific reads from other KIRs, raw genomic sequence reads were mapped onto a complete genomic *Mamu-KIR2DL4* sequence extracted from a bacterial artificial chromosome (bRH-242L13, accession# BX842591) sequenced by Sambrook et al. (2005). All reads that mapped to the 5’UTR and exon one region of *KIR2DL4* contained two mismatches within exon one and three mismatches 3bp upstream of exon one relative to our previous 5’ UTR forward primer. To accommodate this variation, a KIR2DL4-specific primer was designed within a conserved region of the 5’ UTR, shown in Figure 1. We then wanted to investigate potential sequence variation within other KIR loci and so genomic reads were mapped against the *Mamu-KIR3DL1* gene from BAC bRH-242L13 in a similar manner. The majority of animals contained reads with variation in the region targeted by the forward sense PCR primer (data not shown). We also noted the targeted region contained two inframe start codons at amino acid positions one and four, located within the region targeted by the forward primer. Alternative translational start sites are commonly observed in signal sequences (Kochetov et al. 2008). We therefore designed a revised forward primer upstream of these two possible start sites to ensure sequencing of full open reading frames. The revised primer included two degenerate bases to account for observed variation within the 5’ UTRs of *KIR1D* and *KIR3D* transcripts.

**Fig. 1.**
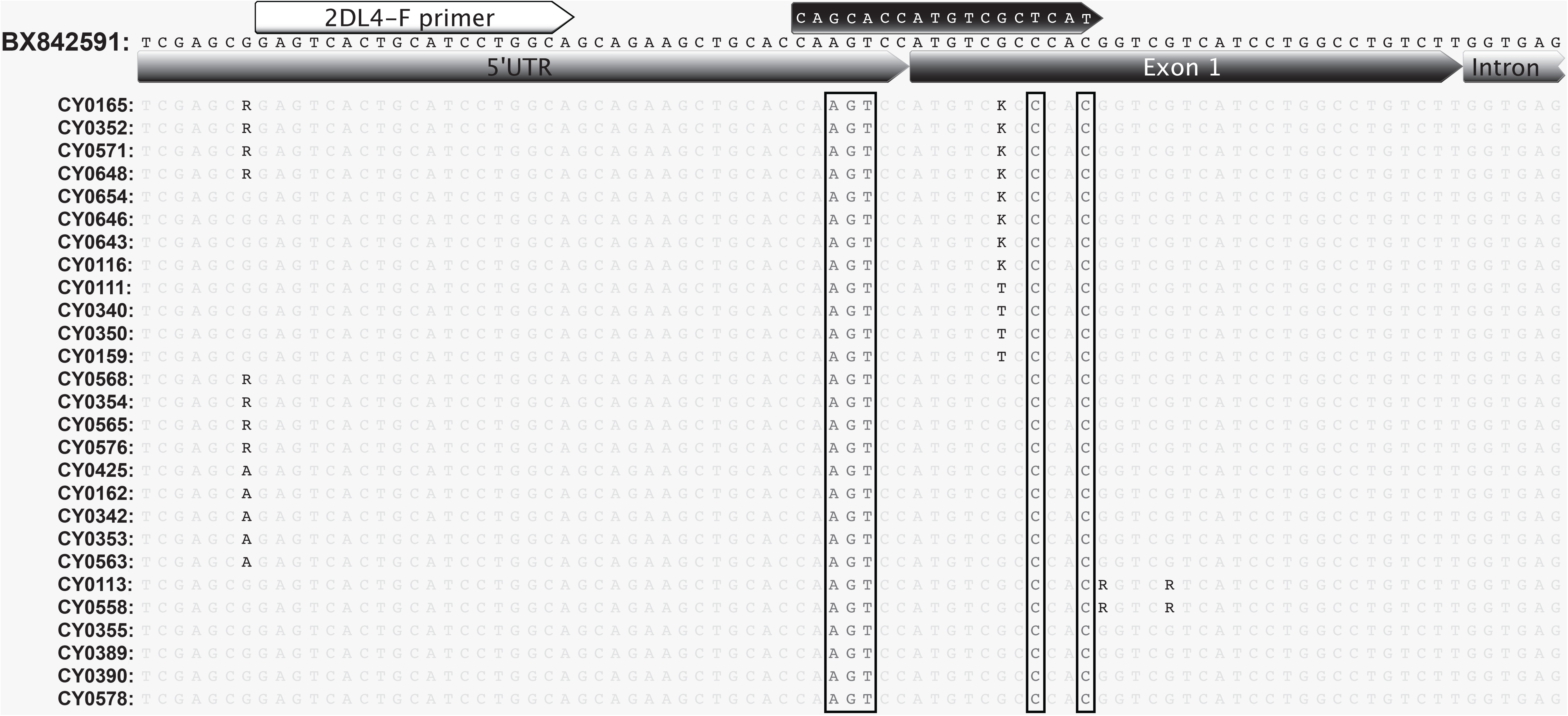
*KIR2DL4* genomic alignment. Genomic reads mapped to BX842591 *KIR2DL4* exon 1. Each sequence represents a consensus of all reads mapped. Degenerate bases are used to represent sites of heterogeneity. Detected genotypes are represented by 4 samples each. Previous forward primer sequence is shown in the black arrow. Mismatches within the previous forward primer targeted region are boxed. The revised primer binding region is shown by the white arrow.

Using revised primer pairs, we generated full-length cDNA amplicons from 30 MCM for PacBio SMRT-CCS analysis. Ten of these 30 individuals had known KIR genotypes determined using microsatellite analysis from our previous study (Supplementary Table 1). These ten animals were selected to represent all eight KIR haplotypes in the MCM population by one or more heterozygous animals; individuals homozygous for the four most common KIR haplotypes were available for this analysis. We successfully confirmed 37 of the 40 previously described *Mafa-KIR* transcripts in MCM, adding 5’ extensions to create full-length open reading frames for 35 of these sequences. The three transcripts that were not detected (EU419100, EU419107, and EU419118) could have been missed due to a variety of reasons in this analysis. EU419107 is the only reported *KIR3DL20* allele from our previous study and is presumably the most centromeric locus within the KIR region. Because of the differences in sequence motifs observed between centromeric and telomeric KIR genes, it is possible we were unable to resequence EU419107 due to mismatches within primer binding sites. EU419118 *(KIR3DL7)* and EU419100 *(KIR3DL2)* are associated with the less common haplotypes K4 and K5, respectively and may have been underrepresented within the pooled library. We suspect that sequencing more MCM may resolve some of the missed transcripts. In addition to the 37 confirmed transcripts, we discovered nine novel KIR transcript sequences plus five additional novel splice variants (Table 1).

**Table 1.** Accession numbers for PacBio-identified sequences

| Haplotype | Gene | Accession | Amino Acids | Representative animals | Prev Accession |
| --- | --- | --- | --- | --- | --- |
| K1 | KIR1D | KX892645 | 279 | CY0161,CY0327 | EU419110 |
| K1K7 | KIR2DL4 | KX892646 | 375 | CY0161,CY0355 | EU419111 |
| K1K7 | KIR2DL4-SV | KX892647 | 277 | CY0563 | EU419111 |
| K1 | KIR3DS | KX892648 | 374 | CY0161,CY0327 | EU419113 |
| K1 | KIR3DL7 | KX892649 | 447 | CY0161,CY0327 | EU419109 |
| K1 | KIR3DL11 | KX892650 | 447 | CY0161,CY0327 | EU419101 |
| K1 | KIR3DL11-SV | KX892651 | 349 | CY0161,CY0327 | EU419102 |
| K2 | KIR1D | KX892652 | 279 | CY0166,CY0568 | EU419124 |
| K2K4K5 | KIR2DL4 | KX892653 | 375 | CY0166,CY0570,CY0390 | EU419114 |
| K2 | KIR3DL2 | KX892654 | 444 | CY0166,CY0353 | EU419090 |
| K2 | KIR3DL2-SV | KX892655 | 394 | CY0425,CY0568 | EU419091 |
| K2 | KIR3DL2 | KX892656 | 444 | CY0166,CY0353 | EU419083 |
| K2 | KIR3DS | KX892657 | 343 | CY0166,CY0353 |  |
| K2 | KIR3DS | KX892658 | 386 | CY0166,CY0568 | EU419092 |
| K2 | KIR3DS | KX892659 | 343 | CY0166,CY0568 | EU419079 |
| K2 | KIR3DL1 | KX892660 | 447 | CY0166,CY0568 | EU419076 |
| K2 | KIR3DL7 | KX892661 | 461 | CY0163 | EU419078 |
| K2 | KIR3DL7 | KX892662 | 447 | CY0166,CY0568 | EU419089 |
| K3K4 | KIR1D | KX892663 | 176 | CY0111,CY0570 | EU419120 |
| K3K4 | KIR1D-SV | KX892664 | 210 | CY0111,CY0570 |  |
| K3 | KIR2DL4 | KX892665 | 375 | CY0111,CY0390 |  |
| K3 | KIR2DL4-SV | KX892666 | 280 | CY0111,CY0390 |  |
| K3 | KIR3DL2 | KX892667 | 444 | CY0111,CY0390 | EU419123 |
| K3 | KIR3DS | KX892668 | 374 | CY0111,CY0390 |  |
| K3 | KIR3DL11 | KX892669 | 447 | CY0111,CY0390 | EU419119 |
| K4 | KIR3DL2 | KX892670 | 444 | CY0570,CY0332 | EU688993 |
| K4 | KIR3DL2-SV | KX892671 | 394 | CY0570,CY0332 |  |
| K4 | KIR3DL2-SV | KX892672 | 435 | CY0570,CY0332 |  |
| K4 | KIR3DS | KX892673 | 374 | CY0570,CY0332 | EU419121 |
| K4 | KIR3DL11 | KX892674 | 447 | CY0570,CY0332 | EU419115 |
| K4 | KIR3DL11-SV | KX892675 | 397 | CY0570,CY0342 | EU419116 |
| K5 | KIR3DL2 | KX892676 | 444 | CY0327,CY0163 | EU419105 |
| K5 | KIR3DL2-SV | KX892677 | 435 | CY0327,CY0163 | EU419106 |
| K5 | KIR3DL2-SV | KX892678 | 394 | CY0327,CY0163 |  |
| K5 | KIR3DS | KX892679 | 374 | CY0327,CY0163 | EU419126 |
| K5 | KIR3DS | KX892680 | 386 | CY0327,CY0163 | EU419097 |
| K5 | KIR3DS | KX892681 | 386 | CY0327,CY0163 | EU419098 |
| K5 | KIR3DL1 | KX892682 | 458 | CY0327,CY0163 | EU419075 |
| K5 | KIR3DL7 | KX892683 | 447 | CY0327,CY0163 | EU419103 |
| K5 | KIR3DL7 | KX892684 | 447 | CY0327,CY0354 | EU419099 |
| K6 | KIR1D | KX892685 | 279 | CY0568 |  |
| K6 | KIR3DL2 | KX892686 | 444 | CY0568,CY0330 |  |
| K6 | KIR3DS | KX892687 | 386 | CY0330 | EU688994 |
| K6 | KIR3DS | KX892688 | 374 | CY0568,CY0330 |  |
| K6 | KIR3DS | KX892689 | 374 | CY0568,CY0330 |  |
| K6 | KIR3DL11 | KX892690 | 447 | CY0568,CY0330 | EU688995 |
| K7 | KIR3DL2 | KX892691 | 440 | CY0356,CY0336 | EU419080 |
| K7 | KIR3DL2-SV | KX892692 | 348 | CY0356,CY0336 | EU419081 |
| K7 | KIR3DS | KX892693 | 374 | CY0356,CY0336 | EU419086 |
| K7 | KIR3DS | KX892694 | 374 | CY0356,CY0336 |  |
| K7 | KIR3DL7 | KX892695 | 447 | CY0356,CY0336 | EU419087 |
| K7 | KIR3DL7-SV | KX892696 | 349 | CY0356,CY0336 | EU419088 |
| K7 | KIR3DL11 | KX892697 | 447 | CY0356,CY0336 | EU419093 |
| K8 | KIR1D | KX892698 | 312 | CY0322,CY0558 |  |
| K8 | KIR3DS | KX892699 | 447 | CY0332 | EU419122 |
| K8 | KIR3DL7 | KX892700 | 397 | CY0322,CY0558 | EU419077 |
| K8 | KIR3DL7 | KX892701 | 374 | CY0322,CY0558 | EU419084 |
| K8 | KIR3DL7-SV | KX892702 | 447 | CY0322,CY0558 | EU419085 |
| K8 | KIR3DL11 | KX892703 | 447 | CY0322,CY0558 | EU419094 |
| K8 | KIR3D11-SV | KX892704 | 349 | CY0322,CY0558 | EU419095 |

KIR genotypes for each animal were determined from a total of 28,086 reads (avg 936 reads/animal) that mapped perfectly to a *Mafa-KIR* allele library consisting of known and novel sequences identified in the current study. Figure 2 displays the genotypes of a subset of sequenced individuals representative of the eight KIR haplotypes in the MCM population. All 37 previously described transcripts were detected on the expected KIR haplotypes with the exception of KX892690 *(KIR3DL11)*, which was associated with K4 haplotypes rather than K6 as previously thought. Of the nine novel sequences we identified two *KIR1D* and one *KIR2DL4* allele. To infer the lineage of the remaining novel sequences we constructed *KIR3DL* and *KIR3DS* phylogenetic trees using all of the sequences containing three immunoglobulin domains that were characterized in this study (Supplementary Fig. 1). Based on phylogenetic analysis the novel sequences were characterized as five *KIR3DS* and one *KIR3DL2* transcript (Table 1). Among the newly identified transcripts was a K3-associated *KIR3DS* (KX892668). No *KIR3DS* alleles were detected for this haplotype in our previous study (Bimber et al. 2008). This establishes *KIR3DS* as the first gene present on all 8 MCM haplotypes. In addition, we discovered new haplotype associations for two previously identified *KIR2DL4* (KX892646, KX892653) and one *KIR1D* (KX892663) transcript. KX892646 *(KIR2DL4)* was associated with both K1 and K7 haplotypes; KX892653 *(KIR2DL4)* was associated with K2, K4 and K5 haplotypes; and KX892663 *(KIR1D)* was associated with K3 and K4 haplotypes. Similar observations have been reported in rhesus macaques where identical *Mamu-KIR1D* and *Mamu-KIR2DL4* alleles are shared between haplotypes (Blokhuis et al. 2010). The combined novel transcripts and haplotype associations have added transcripts to seven of the eight haplotypes. The range of transcripts per haplotype increased to between five and nine.

**Fig. 2.**
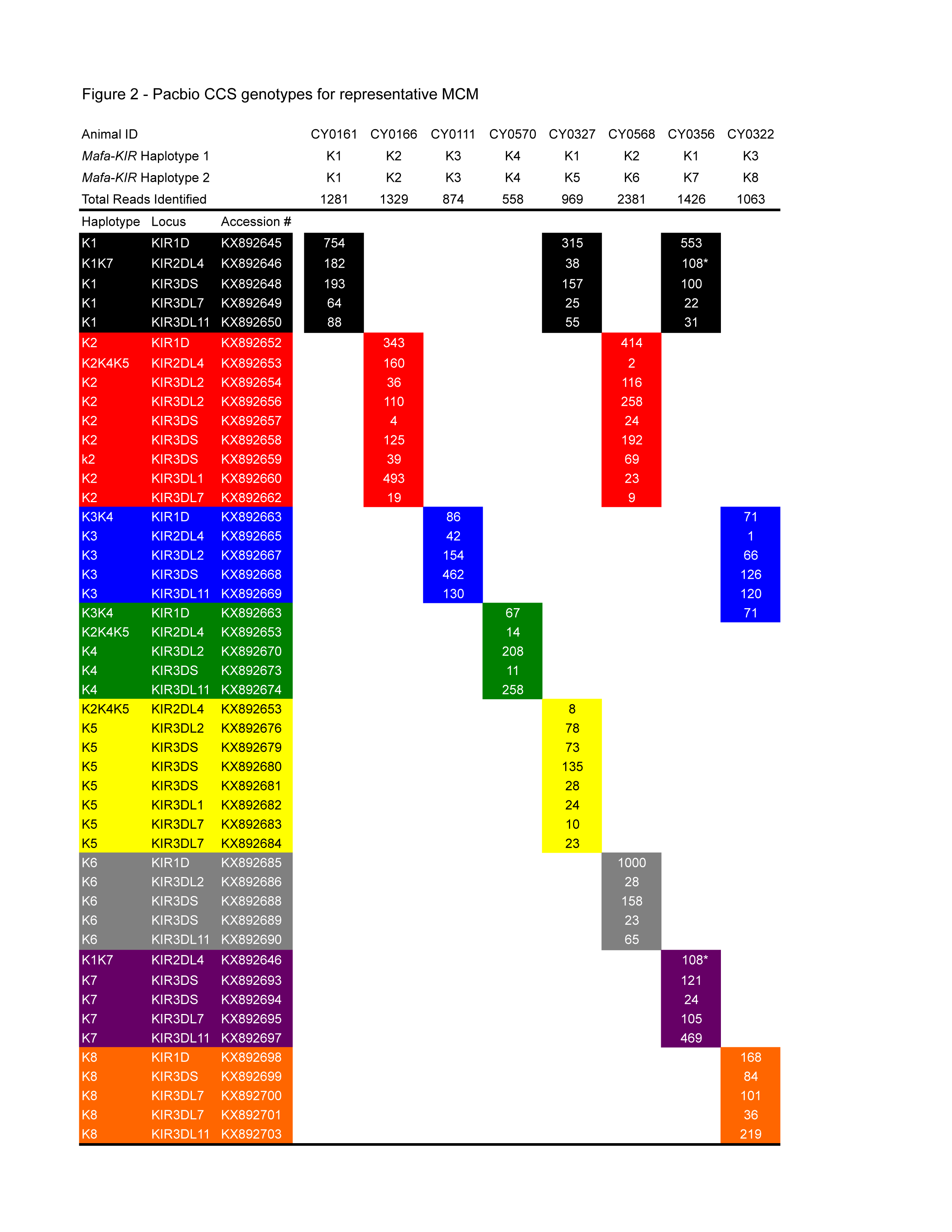
PacBio CCS genotypes for representative MCM. The number of identical CCS reads for each transcript is presented within cells. Columns represents CCS reads detected for individuals. Haplotype associations are designated by color. * denotes sequence read counts that are shared between 2 haplotypes.

### Genotyping from WGS and MES datasets

In previous studies, we and others (Vallender 2011) have used human exome probe arrays to capture and sequence exons from the rhesus and cynomolgus macaques. However, using probes designed for the human genome (VCRome2.1, total length of capture probes 35.25 Mb) to capture the rhesus macaque exome failed to produce sufficient read coverage for 12% of exons in the rheMac2 annotated rhesus genome assembly (Bainbridge et al. 2011). Therefore, to improve overall coverage of macaque exomes and simultaneously determine macaque MHC class I and class II genotypes, we designed additional capture probes targeting rhesus exons that were not covered sufficiently using VCRome2.1 capture reagents (Supplementary Table 3). A series of tests comparing the results for macaque exome sequencing using the combined VCRome2.1 plus “spike-in” reagents against whole genome sequencing results from the same animals showed that a ratio of 1:2.5 for VCRome2.1-to-“spike-in” provided superior overall coverage results (Supplementary Tables 4 and 5).

With a full-length *Mafa-KIR* alleles database established, we evaluated the feasibility of genotyping KIRs directly from MES reads compared to reads generated using WGS. All 30 PacBio-sequenced animals were genotyped using their respective MES (n=23) and WGS (n=7) datasets. 100bp paired end reads were mapped to a library of *Mafa-KIR* D0, D1, and D2 exon sequences. Genotypes for the 30 PacBio-sequenced animals were determined by requiring complete coverage of D0, D1, and D2 for *KIR3DL/KIR3DS* alleles; D0 and D2 for *KIR2DL4* alleles; and D1 for *KIR1D* alleles. KIR alleles detected per haplotype for representative MCM are shown in Figure 3. In total, we were able to detect 42 of the 45 PacBio-identified transcripts. Sequence depths between WGS and MES were comparable. Interestingly, all three previously identified alleles that were absent within our PacBio sequencing experiment (EU419100, EU419107, and EU419118) were successfully mapped from genomic reads (Supplementary Table 6). These KIR alleles may either represent KIR alleles with sequence variants in the PacBio primer binding sites or transcripts with low expression levels in the animals evaluated in this study.

**Fig. 3.**
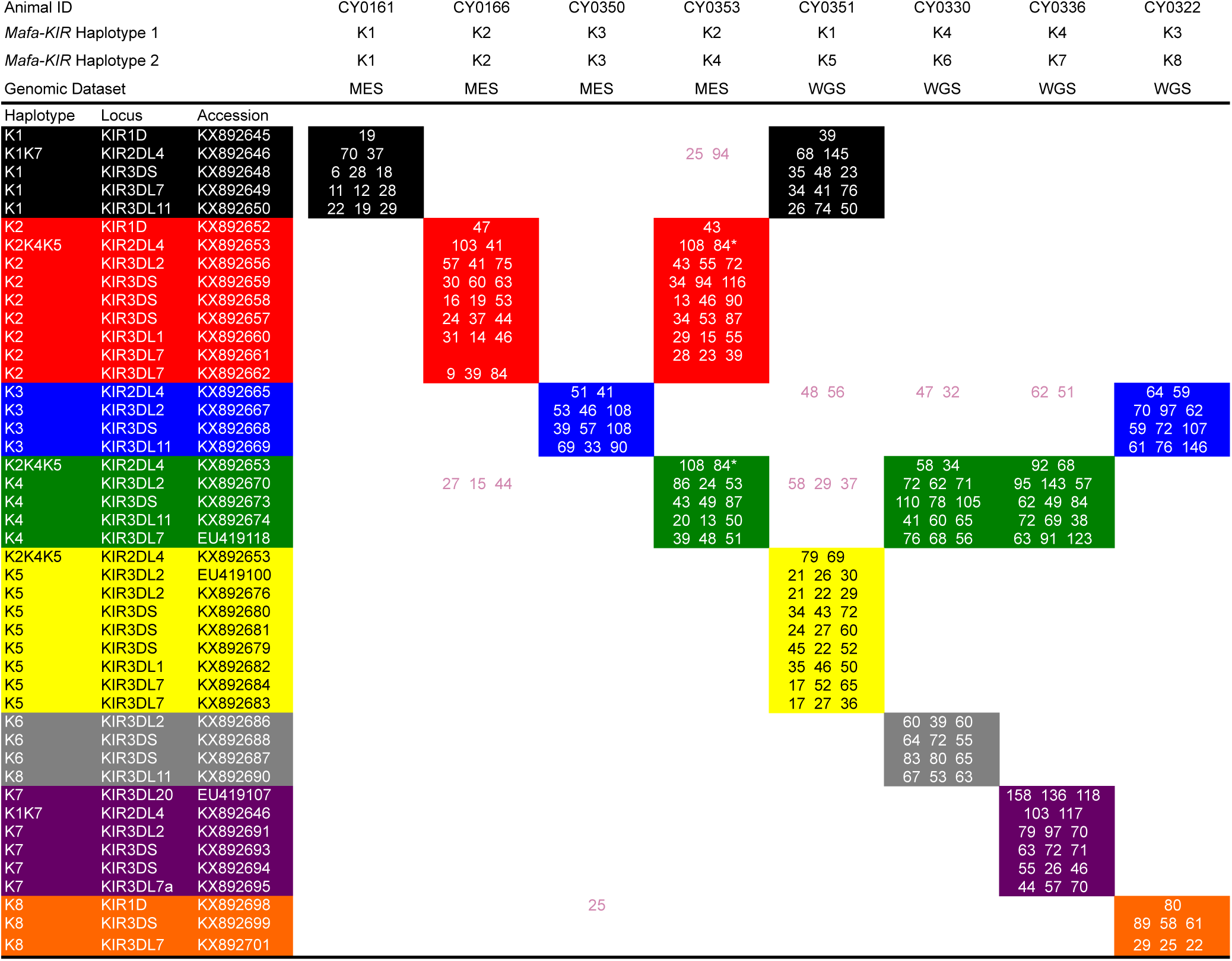
WGS/MES genotypes for representative MCM. The number of sequence reads mapped to representative exons are presented within a single cell. Three numbers are given for *KIR3DS/KIR3DL* alleles, two for *KIR2DL4* alleles, and one for *KIR1D* alleles. Haplotype associations are designated by color. * denotes sequence read counts that are shared between 2 haplotypes.

## DISCUSSION

Previously, we and others have developed PacBio CCS methods for full length MHC class I allele discovery and genotyping (Westbrook et al. 2015) (Karl et al. under review: http://biorxiv.org/content/early/2016/11/02/084947). Here we adapted this framework to full-length allele discovery and genotyping for KIRs. Together, these methods allow for characterization of macaque KIR/MHC genetics without the ambiguity of genotyping based on sequences of partial gene fragments. In addition, by utilizing genomic sequence data we were able to accommodate multiple sites of variation when designing primers, increasing our sensitivity of detection. This approach allowed us to detect nine novel transcripts, five novel splice variants, and four novel haplotype associations within MCM where KIR diversity was thought to be completely characterized. Our reported findings provide a more comprehensive perspective of the genetics of KIRs within MCM.

PacBio analyses revealed a novel *KIR3DS* transcript associated with K3 haplotypes; no *KIR3DS* alleles were detected for this haplotype in our previous study (Bimber et al. 2008). Thus, at least one *KIR3DS* gene is present on all eight KIR haplotypes in MCM. It is therefore possible that a *Mafa-KIR3DS* lineage represents a framework gene within MCM. *KIR3DS* (formerly *KIR3DH)* in macaques describes a class of receptors that are evolutionarily distinct from human *KIR3DS*, most likely arising from a crossover event between a *KIR3DL* and *KIR2DL4* within an ancestral macaque (Hershberger et al. 2001). Like *KIR2DL4, KIR3DS* genes contain a charged arginine residue within their transmembrane domains. *KIR3DS* also contain a large deletion resulting in early termination before the immunoreceptor tyrosine-based inhibitory motifs (ITIMs) encoded by KIR2DL4 and KIR3DL cytoplasmic tails (Hershberger et al. 2001). Therefore, KIR3DS are thought to be the only activating receptors in macaques. Several haplotypes that lack *KIR3DS* have been reported in rhesus macaques including the single *Mamu-KIR* haplotype that was completely sequenced from two overlapping BACs (Blokhuis et al. 2011; Kruse et al. 2010; Moreland et al. 2011, Sambrook et al. 2005). However, *Mamu-KIR* genotyping studies from a variety of groups have shown that virtually all rhesus encode at least one *KIR3DS* activating receptor (Blokhuis et al. 2009; Blokhuis et al. 2010; Hellmann et al. 2011; Hershberger et al. 2001; Kruse et al. 2010; Moreland et al. 2011). In all of the sequencing literature from the past decade we could find only a handful of rhesus macaques reported with no *Mamu-KIR3DS* (Blokhuis et al. 2011). While it is possible that rare macaque haplotypes lack *KIR3DS* it seems likely that *KIR3DS* variants may be present in these animals that were not detected by the various genotyping assays used in these earlier studies. Taken with our findings that activating KIR are expressed in all MCM, it is tempting to speculate that activating KIR provide some a selective advantage.

A growing amount of evidence suggesting the role of particular activating *Mamu-KIR3DS* alleles in influencing plasma viral loads in SIV-infected rhesus macaques has surfaced from associative studies (Albrecht et al. 2014; Chaichompoo et al. 2010). Similarly, the copy number of *Mamu-KIR3DS* genes is speculated to influence peak viral loads following SIVmac251 infection in Indian rhesus macaques. Hellmann et al. (2011) showed that increased copies of *Mamu-KIR3DS* were inversely correlated with peak viral loads, though it should be noted that the association was seen only in the rhesus that lack protective *Mamu-A^*001^* and were homozygous for restrictive *TRIM5* alleles (Hellmann et al. 2011). The methods described in this study could aid in untangling the biological mechanisms of activating KIRs in future studies by providing unambiguous genotyping down to the allelic identity. Furthermore, copy number variation of activating *KIR3DS* exists across MCM haplotypes. Investigating activating *KIR3DS* copy number variation using MHC/KIR identical cohorts may lead to novel insights into these complex associations.

*KIR2DL4* is considered a framework gene in humans and is thought to play a role in recognition of fetal tissues by the maternal immune system through binding nonclassical MHC-G molecules (Ponte et al. 1999). Data by Yan et al. (2007) has shown that *KIR2DL4* surface expression in uterine NK was significantly lower in women with recurrent spontaneous abortion than in normal early pregnancy woman. Our study found that the two previously identified *KIR2DL4* sequences are shared among multiple haplotypes. In addition, a novel K3-associated *KIR2DL4* was characterized making six of the eight haplotypes *KIR2DL4* positive. When we genotyped animals using WGS/MES reads, all 30 samples had complete coverage of both *KIR2DL4* exons for at least one of the three alleles (data not shown). This was not seen in our PacBio cohort as not every sequenced animal contained *KIR2DL4* CCS reads. It is possible that WGS/MES *KIR2DL4* reads are from pseudogenes within these animals; however, it is also possible that the absence of *KIR2DL4* CCS reads was due to inefficient PCR amplification or low transcript abundance. The latter explanation is interesting when considering that all 62 genomic datasets contained reads spanning the first exon of *KIR2DL4* including an in-frame start codon when mapped against the *Mamu-KIR2DL4* contig extracted from the genomic haplotype (Fig. 1). Furthermore, seven unique *KIR2DL4* 5’UTR sequences were identified within the genomic datasets but only three unique *KIR2DL4* transcripts were identified with PacBio sequencing. It’s known that the relative abundance of KIR transcripts fluctuates over the course of infection due to selective expansion of NK cell populations (reviewed in Maras et al. 2014). Any of these factors may have contributed to the varying levels of CCS read depth we observed across our Pacbio cohort. Interestingly, six animals were positive for all three *KIR2DL4* respectively when genotyped using WGS/MES data. This could be a byproduct of ambiguous mapping of short reads onto larger exon contigs. However, this observation is consistent in rhesus macaques where Hellmann et al. (2013) found *Mamu-KIR2DL4* copy numbers varied between one and three per cell. Understanding *KIR2DL4* presence/absence and copy number variation requires sequencing of more animals to identify homozygous individuals with less common KIR haplotypes.

In this study we also demonstrated an approach to KIR genotyping directly from WGS/MES reads using a reference library of D0, D1, and D2 exon sequences derived from full-length transcripts. Though we successfully detected 42 total alleles, we were unsuccessful in detecting seven PacBio-identified alleles using WGS/MES genotyping (Supplementary Table 6). This is most likely due to the strict constraints of the WGS/MES genotyping algorithm. For example, CY0322 was identified as a K3/K8 heterozygote (Supplementary Table 1). One hundred and one reads for a K8-associated *KIR3DL7,* KX892700, were detected using PacBio amplicon sequencing (not shown). This allele wasn’t detected from the same animal using mapped WGS reads. Viewing the per-exon coverage for KX892700 reveals complete coverage for D2, but incomplete coverage for D0 and D1. Similarly, a K2-associated *KIR3DL7,* KX892661, appears to be absent in MES data for CY0166 who is a K2/K2 homozygous animal. Again, viewing the per exon coverage for CY0166 reveals that complete D0 and D2 coverage exists for KX892661, but D1 has incomplete coverage and thus the allele was not scored as positive. One solution to this problem may be to slightly decrease the coverage required to call an allele or shorten the length of reference contigs that reads are mapped to. However, this could potentially complicate genotyping in instances where alleles differ by only a few nucleotides. Improved sequencing platforms such as the Illumina HiSeq X, which generate greater amounts of data per lane at lower cost that earlier platforms, may also help by providing greater read depth across KIR genes at no additional cost. Another solution may be to spike-in KIR probes into the Rhexome1.0 just as we have with the MHC in order to increase read depth. Though the Rhexome1.0 was able to capture a significant amount of KIR sequences without spike-in probes, greater read depth may help increase confidence when determining genotypes. Because of the restricted genetic diversity of MCM, spike-in KIR probes may not be necessary as genotypes can be inferred based on the full-length sequencing studies we have performed to define the extent of KIR diversity within the population. Inference will most likely fall short once more genetically diverse populations are considered such as Indian rhesus macaques. Regardless, applying this genotyping approach to other species will require full-length allele libraries to be established for each population.

This genotyping approach has given us a framework for interpreting complex, duplicated families of genes from genomic data, a task that has not been possible using reference guided alignment given the current genome assemblies for nonhuman primates. Currently we are working to expand this methodology to other complex immune gene families such as MHC class I, class II and Fc gamma receptors. This would eventually allow genotyping of a variety of immune loci directly from a single MES experiment.

## ACKNOWLEDGEMENTS

This research was supported by contracts HHSN272201600007C and HHSN272201100013C from the National Institute of Allergy and Infectious Diseases of the National Institutes of Health, and was conducted at a facility constructed with support from the Research Facilities Improvement Program (RR15459-01, RR20141-01).

## SUPPLEMENTARY MATERIALS

**Supplementary Fig. 1** — *Mafa-KIR3DL* and *Mafa −KIR3DS* phylogenetic trees Phylogenetic analysis of *KIR3D* sequences identified with pacbio sequencing. Black diamonds indicate novel sequences. A, The tree contains *KIR3DL* sequences. Black bars represent *KIR3DL* lineage groups. B, The tree contains KIR3DS sequences.

**Supplementary Table 1** — Pacbio CCS cohort

**Supplementary Table 2** — WGS/WES/MES cohort

**Supplementary Table 3** — Rhexome1.0 target capture components

**Supplementary Table 4** — Coverage from VCRome2.1-to-“spike-in” mixing ratios for a single rhesus macaque.

**Supplementary Table 5** — MES target capture coverage for 15 rhesus macaques

**Supplementary Table 6** — Comparison of sequences identified by Pacbio CCS versus WGS/MES

